# *Keratinocyte-Associated Protein 3* is a novel gene for adiposity with differential effects in males and females

**DOI:** 10.1101/2022.02.20.481201

**Authors:** Alexandria M Szalanczy, Emily Goff, Osborne Seshie, Aaron Deal, Michael Grzybowski, Jason Klotz, Chia-Chi C Key, Aron M Geurts, Leah C Solberg Woods

**Author notes:** **Contact Info:** Leah C Solberg Woods, Department of Internal Medicine, School of Medicine, Wake Forest University, Winston Salem, North Carolina, USA. **Author Contributions** AMS designed the study, conducted experiments, ran statistical analysis, analyzed results, created figures, and wrote manuscript. LSW designed study, oversaw experimental work, and edited manuscript. EG, OS, AD, and CCK assisted with experiments. MG, JK, and AMG created rat knock-out model. All authors approved final version of the manuscript.

## Abstract

**Objective:** Despite the obesity crisis in the United States, the underlying genetics are poorly understood. Our lab previously identified *Keratinocyte-associated protein 3, Krtcap3,* as a candidate gene for adiposity where increased expression of *Krtcap3* correlated with decreased fat mass. Here we seek to confirm that *Krtcap3* expression affects adiposity traits.

**Methods:** We developed an *in vivo* whole-body *Krtcap3* knock-out (KO) rat model. Wild-type (WT) and KO rats were placed onto a high-fat or low-fat diet at six weeks of age and were maintained on diet for 13 weeks, followed by assessments of metabolic health. We hypothesized that *Krtcap3*-KO rats will have increased adiposity and a worsened metabolic phenotype relative to WT.

**Results:** We found that KO male and female rats have significantly increased body weight versus WT. KO females ate more, had more fat mass, but were also more insulin sensitive than WT. Alternatively, KO males weighed more and were more insulin resistant than WT, with no differences in eating or fat mass.

**Conclusions:** This study validates *Krtcap3* in body weight regulation and demonstrates sex-specific effects on food intake, adiposity, and insulin sensitivity. Future studies will investigate how *Krtcap3* is acting and seek to better understand these sex differences.

**Study Importance Questions:** What is already known about this subject?

- Over 900 low-risk, common genetic variants for BMI have been identified, but these still only explain a fraction of the heritability and many of the underlying causal genes remain unknown
- *Krtcap3* has been identified as a candidate gene for obesity in both rats and humans, but no verification or functional studies have been done

What are the new findings in your manuscript?

- Identified *Krtcap3* as a novel gene that impacts feeding behavior and adiposity in female rats
- Determined that *Krtcap3* impacts insulin sensitivity differentially in male and female rats

How might your results change the direction of research or the focus of clinical practice?

- This work may lead to identification of new pathways that contribute to obesity without metabolic complications, which will advance understanding of the biology of obesity and potentially identify novel drug targets
- This work highlights the need to investigate sex differences in the genetics of obesity

## Introduction

Global obesity rates have nearly tripled in the last 50 years, and in the United States adult obesity rates are expected to reach 50% by 2030 (1, 2). Obesity is influenced by both environment and genetics (3, 4, 5), but the genetic architecture underlying the disease is poorly understood. Although many genetic loci have been identified in human genome wide association studies (GWAS), many of the causal genes are unknown. Additionally, known loci explain only a small portion of heritability, revealing significant gaps in knowledge (4, 5, 6).

Our laboratory has used the outbred heterogeneous stock (HS) rat as a complementary method for genetic mapping of obesity traits (7, 8). HS rats are outbred from eight inbred founder strains (9), allowing genetic fine-mapping to Mb intervals (10). Previously, we used mediation analysis to identify *Keratinocyte-associated protein 3* (*Krtcap3*) as a candidate gene for visceral fat mass within a quantitative trait locus (QTL) on rat chromosome 6 (7). Liver *Krtcap3* expression was negatively correlated with fat mass, suggesting a protective role against obesity. Another group also found that *Krtcap3* is a potential pleiotropic gene for obesity, type 2 diabetes (T2D), and dyslipidemia in humans (11), indicating translational relevance. That said, this gene lacks validation.

To determine the role of *Krtcap3* on adiposity, we knocked-out *Krtcap3* expression in an *in vivo* WKY rat model, as the WKY strain’s haplotype conferred susceptibility high liver *Krtcap3* expression and decreased fat mass at the chromosome 6 QTL (7). Wild-type (WT) and knock-out (KO) rats were placed on either a low-fat (LFD) or high-fat diet (HFD) and multiple metabolic traits were measured. We show that *Krtcap3-*KO rats have increased body weight in both sexes, but increased food intake, fat mass, and insulin sensitivity only in female KO rats. This work validates the role of *Krtcap3* in adiposity, and demonstrates sex-specific differences on its impact on food intake, adiposity, and insulin sensitivity.

## Methods

### Animals

We used CRISPR-Cas9 to develop a *Krtcap3*-KO (WKY-Krtcap3^em3Mcwi^) by deleting two base pairs in exon 2, using the Wistar-Kyoto (WKY/NCrl; RGD_1358112) as the genetic background strain (**Figure 1a**). In addition to conferring susceptibility to decreased fat mass at the chromosome 6 QTL, the WKY strain has naturally high liver *Krtcap3* expression relative to other inbred rat strains, although lower adipose *Krtcap3* expression (**Figure S1**). Primers are given in Table S1. We used real-time quantitative PCR (rt-qPCR) to verify significantly decreased *Krtcap3* expression levels in the liver of the *Krtcap3-*KO relative to WT (T_4_=14.44, p=6.67e-5) (**Figure 1b**).

**Figure 1.**
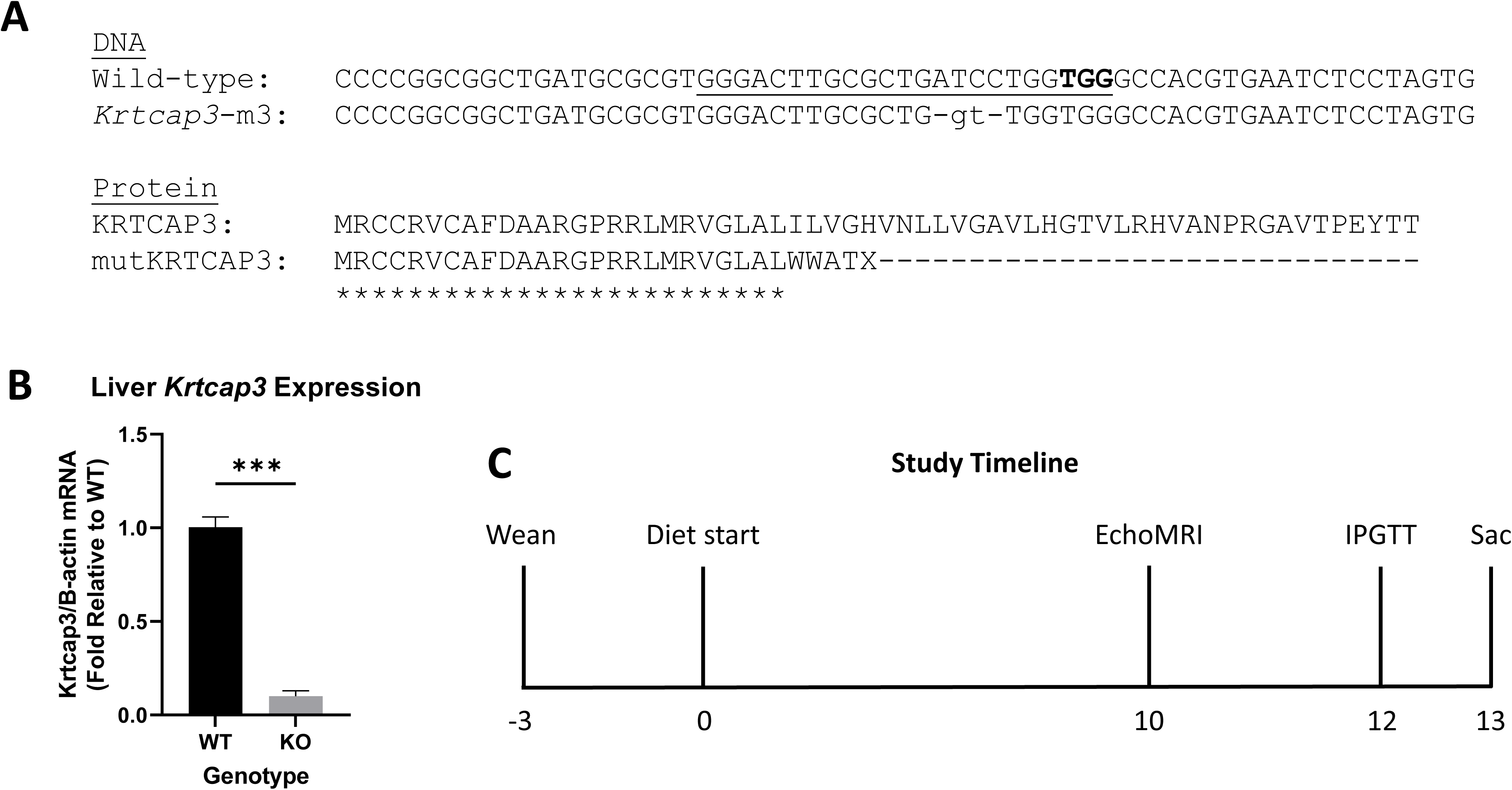
Development and confirmation of *Krtcap3* knock-out (KO) rats and study design. (A) CRISPR-Cas9 mutagenesis was used to target the second exon (underlined sequence, protospacer adjacent motif in bold), resulting in a 4 base-pair deletion and 2-based pair insertion (lowercase). This results in a predicted frameshift after 25 amino acids and premature termination of the normal protein sequence (only the first 60 of 240 amino acids shown). Asterisks indicate identical amino acids. (B) RNA was extracted from liver tissue of wild-type (WT) and KO rats (n=3 per genotype) and *Krtcap3* expression analyzed via quantitative PCR to assess the success of the *Krtcap3*-KO. Fold change was calculated relative to WT *Krtcap3* expression. ***p < 0.001. (C) Timeline outlining study design, with weeks relative to diet start shown. Metabolic phenotyping included EchoMRI analysis (EchoMRI), an intraperitoneal glucose tolerance test (IPGTT), and euthanasia (Sac).

Three heterozygous (HET) breeding pairs were used to establish a breeding colony at Wake Forest University School of Medicine (WFSoM). Rats were housed in standard caging at 22°C in a 12 h light and 12 h dark cycle (dark from 6:00 pm to 6:00 am) at standard temperature and humidity conditions and given *ad libitum* access to water. Breeders were given standard chow diet while experimental rats were placed on experimental diet as described below.

### Study Design

Experimental rats were weaned at three weeks of age and placed two per cage in same-sex, same-genotype cages. At six weeks of age, rats were weighed and began either a HFD (60 %kcal fat; ResearchDiet D12492) or a sucrose-matched LFD (10 %kcal fat; ResearchDiet D12450J) (**Table S2**). Hereafter rats will be referred to by genotype-diet-sex, such that WT males consuming HFD are called WT HFD males. Rats were allowed access to diet *ad libitum* with body weight and food intake recorded weekly. Rats were on diet for 13 weeks, with metabolic phenotyping tests beginning after 10 weeks on diet and euthanasia after 13 weeks on diet, as described (**Figure 1c**). We saved 8-11 rats per genotype-diet-sex group. Due to COVID-19 shut-downs, we were unable to complete the experimental procedure in all rats (**Table S3**). All experiments were performed using a protocol approved by the Institutional Animal Care and Use Committee at WFSoM.

### Metabolic phenotyping

After 10 weeks on diet, rats went through EchoMRI (EchoMRI LLC, Houston, TX) analysis, which precisely measures fat mass and lean mass of live rats. Non-fasted rats were weighed before analysis then scanned for 2 minutes in triplicate.

After 12 weeks on diet, rats were fasted 16 h overnight before being administered an intraperitoneal glucose tolerance test (IPGTT). We collected blood glucose (Contour Next EZ) and serum for subsequent insulin analysis from a tail nick at fasting and 15, 30, 60, and 90 minutes after a 1 mg/kg glucose injection. To calculate a rat’s response to the glucose challenge, the glucose area-under-the-curve (AUC), we used the following equation:

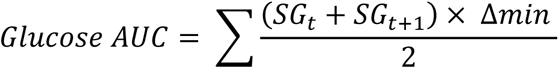

Where serum glucose (SG) was measured at t=0, 15, 30, 60, 90 min.

### Tissue Harvest

After 13 weeks on diet, rats were euthanized via live decapitation. Trunk blood was collected and serum saved and stored for lipid and biochemistry analysis. Fat pad tissues (retroperitoneal ((RetroFat)), gonadal, and omental/mesenteric fat (OmenFat)) and liver were dissected, weighed, and snap-frozen. Pancreas was weighed, saved on ice in acid ethanol, chopped by scissors the same day, and processed 48 hours later and stored at -20 C for subsequent measurement of whole pancreas insulin content (WPIC). Additional sections of liver and RetroFat were saved in 10% neutral-buffered formalin for histology. Due to research facility restrictions at the onset of COVID-19, the tissue harvest protocol had to be shortened, and OmenFat and pancreas for WPIC were not collected from all animals (**Table S3**).

### Insulin

We used ultrasensitive ELISA kits (Alpco Ref # 80-INSRTU-E10) to analyze serum insulin from the IPGTT and WPIC. Insulin response to the IPGTT, insulin AUC, was calculated with the same equation as glucose AUC. WPIC was normalized to pancreas weight. Homeostatic model assessment for insulin resistance (HOMA-IR) was used to approximate insulin resistance and was calculated by the formula (12):

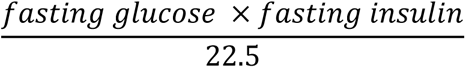

### Serum and liver analyses

Trunk blood was collected at the time of euthanasia, and serum from each rat saved. Undiluted serum from each rat was sent to IDEXX BioAnalytics for analysis on a custom chemistry panel (62761), including measurements of triglycerides and total cholesterol. Lipids were also extracted from liver tissue to measure triglyceride (Wako Diagnostics L-Type TG M) (13) and cholesterol (Pointe Scientific Cholesterol Liquid Reagents).

### Histology

We chose representative WT and KO samples from the HFD female group, where we saw the largest phenotypic difference in RetroFat (n=5 per genotype). The Wake Forest Comparative Pathology Lab stained RetroFat samples for hematoxylin and eosin. Slides were electronically scanned by Wake Forest Virtual Microscopy Core, then analyzed with Visiopharm software (Hoersholm, Denmark). The area of 1000 adipocytes per slide (three slides per rat) were measured within at least three regions of interest per slide and plotted in a frequency distribution assessing adipocyte size. Only adipocytes with areas between 300 μm^2^ and 15,000 μm^2^ were considered.

### Statistical analysis

All data were analyzed in R (1.4.1103) with males and females analyzed separately. Outliers were assessed separately for each of the genotype-diet-sex groups as described in Supplementary Methods. Data were transformed to reflect a normal distribution (**Table S4**).

Wean weight and six-week body weight between WT and KO rats were assessed with a Student’s t-test. Growth curves and cage food intake were assessed with a two-way repeated measures ANOVA, where the effect of genotype and diet were examined over time. If we saw a significant three-way interaction, data were separated by diet to analyze the effect of genotype over time. With the exception of histology data, all other data were analyzed by a two-way ANOVA, where the factors were genotype and diet. If there was a significant interaction, the analysis was split by diet condition and a Student’s t-test assessed the effect of genotype.

We assessed the frequency distributions of adipocyte size between WT and KO rats by sorting adipocyte areas into bins from 0 to 15,000 μm^2^ in increments of 500 μm^2^. The frequency of each bin was calculated and the frequency distributions plotted. Data were analyzed by a two-way ANOVA, with the factors genotype and bin size.

## Results

### Male and female KO rats are heavier than WT at six weeks of age

Although there were no differences in wean weight between WT and KO males or females (**Figure 2a**), by six weeks of age KO males and females were heavier than WT controls (T_34_=2.51, p=0.0085; T_35_=2.43, p=0.010, respectively) (**Figure 2b**). At six weeks of age, when rats began experimental diet, there were no significant differences in body weight between diet groups for either sex or genotype.

**Figure 2.**
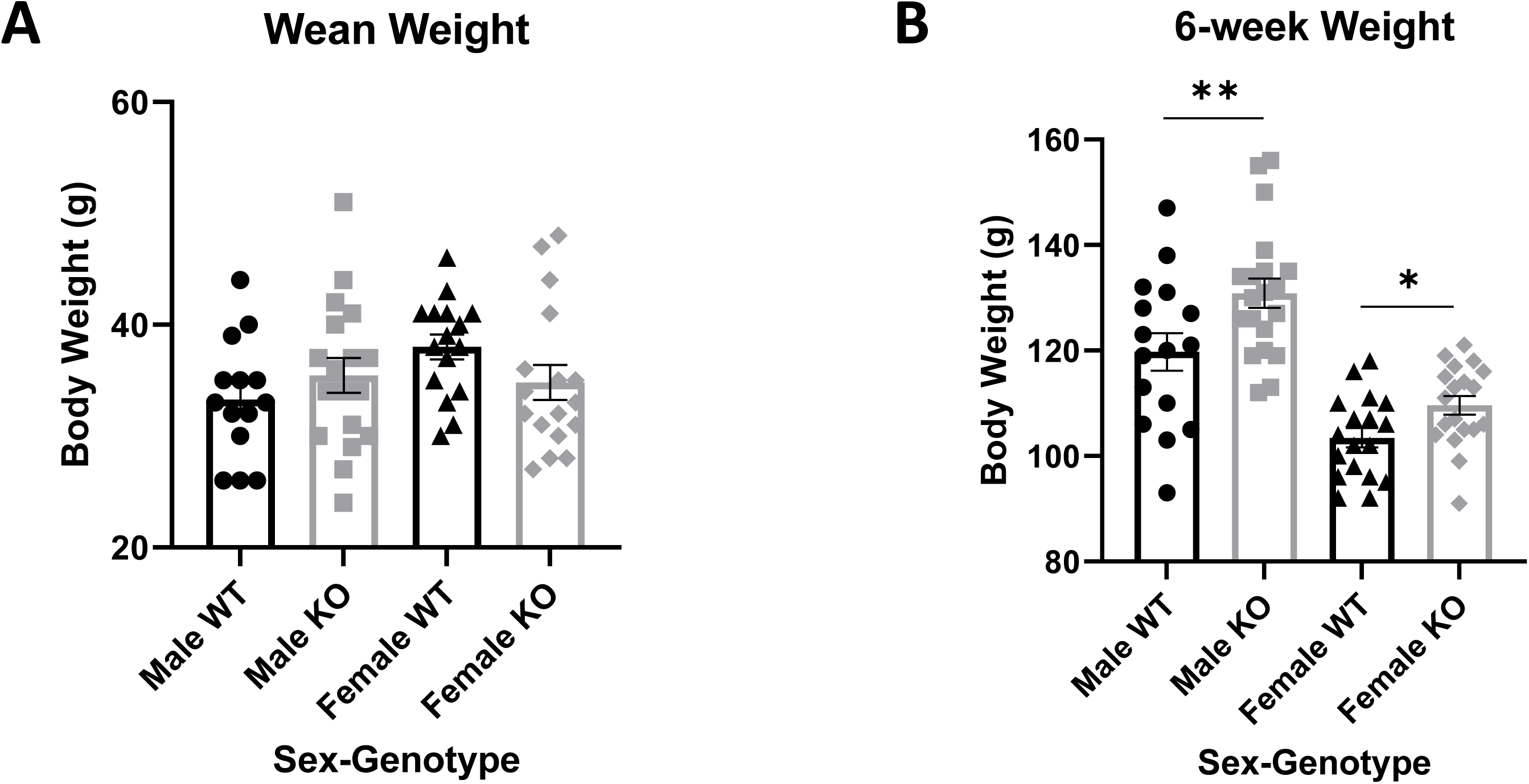
Body weight in wild-type (WT, black) and *Krtcap3* knock-out (KO, gray) rats prior to diet start. (A) At time of wean at three weeks of age, there were no differences in body weight between WT males (circle) and KO males (square), nor between WT females (triangle) and KO females (diamond). By diet start at six weeks of age, however, (B) KO rats of both sexes were significantly heavier than WT controls. *p < 0.05, **p < 0.01.

### Male KO rats are heavier than WT with no difference in food consumption

Over 13 weeks on diet, KO males were heavier than WT males (F_1, 32_=4.44, p=0.043; **Figure 3a**) and HFD males were heavier than LFD (F_1, 32_=4.68, p=0.038). There were no differences in food intake by genotype (**Figure 3b**), but LFD males ate significantly more than HFD males (F_1, 11_=120.18, p=2.93e-7), while still consuming fewer kcal than HFD males (**Figure S2a**).

**Figure 3.**
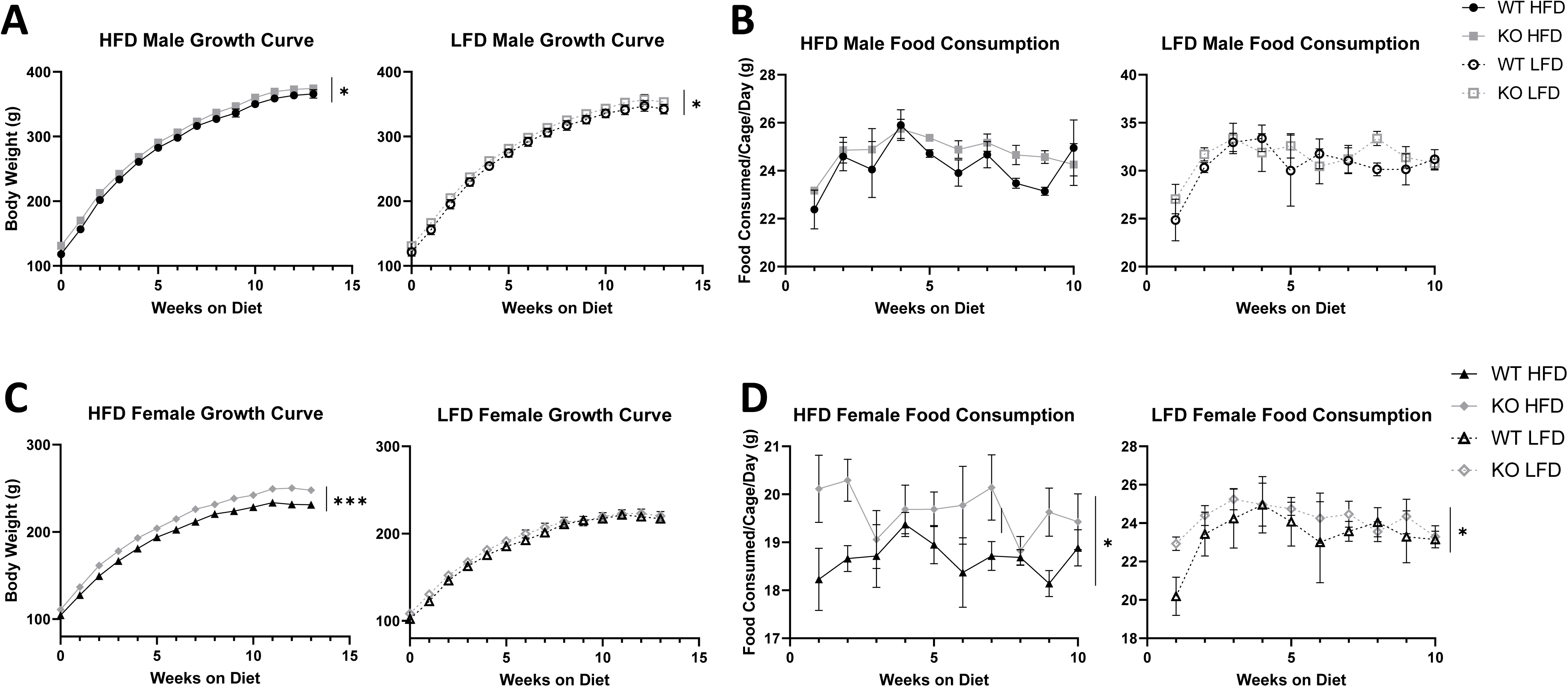
Growth curves and weekly cage food intake in wild-type (WT; black) and *Krtcap3* knock-out (KO; gray) rats. Rats were weighed weekly and cage food intake was recorded. Male data are presented on top, and female data presented on bottom. To highlight genotype-driven differences, figures are separated by diet, high-fat diet (HFD; filled) and low-fat diet (LFD; empty). (A) KO males (square) were consistently heavier than WT males (circle) on both diets. Significance represents the main effect of genotype across both diets. *p < 0.05 for a main effect of genotype across both diets. (B) There were no differences in eating behavior between WT and KO males on either diet, but note the scale difference demonstrating that LFD males consume more food than HFD males. (C) In females, there was a main effect of diet as well as a three-way interaction between genotype, diet, and time. When data was analyzed separately by diet condition, KO females (diamond) on a HFD weighed more than WT (triangle) counterparts, with no difference in weight between WT and KO LFD females. ***p < 0.001 for the effect of genotype in the HFD condition. (D) There was a main effect of genotype on cage food intake for females with no interaction between genotype, diet, and time, where KO females ate more food than WT females. Similar to males, LFD females ate more than HFD females (note scale differences). *p < 0.05 for a main effect of genotype across both diets. Diet effects are reported in the “Results” section.

### Female KO rats are heavier and eat more than WT

Over the course of the study, KO females were significantly heavier than WT females (F_1, 32_=11.54, p=0.0018; **Figure 3c**) and HFD females were heavier than LFD (F_1, 32_=16.22, p=3.2e-4). Visually there is a larger effect in KO HFD females relative to all other groups, which is supported by a significant three-way interaction between genotype, diet, and time (F_13, 416_=2.69, p=0.0012). When separated by diet, KO HFD females weighed significantly more than WT (F_1, 17_=18.44, p=4.9e-4), with no differences by genotype in LFD females. KO females also ate more than WT (F_1, 13_=7.54, p=0.017; **Figure 3d**) and LFD females ate more than HFD (F_1, 13_=100.82, p=1.72e-7), with no interaction between genotype and diet. As with males, HFD females still consumed more kcal than LFD (**Figure S2b**).

### Male KO rats exhibit no difference in fat mass relative to WT, despite increased body weight

After 10 weeks on diet at EchoMRI analysis, there were no differences between WT and KO male total fat mass (**Figure 4a**), but HFD males had more fat mass than LFD (F_1, 33_=54.53, p=1.76e-8). KO males had a trend toward greater lean mass than WT (F_1, 33_=3.14, p=0.086), with no differences in lean mass by diet (**Figure S3a**). At study completion, KO male rats were heavier than WT (F_1, 32_=4.36, p=0.045; **Figure 4b**), and HFD males were heavier than LFD (F_1, 32_=16.36, p=3.1e-4), with no interactions between genotype and diet. Despite the differences in body weight, and consistent with EchoMRI analysis, there were no genotype-driven differences in RetroFat (**Figure 4c**), although HFD males had greater RetroFat than LFD (F_1, 32_=52.08, p=3.39e-8). Likewise, there was no difference in gonadal fat between WT and KO males (**Figure 4d**), but HFD males had increased gonadal fat relative to LFD (F_1, 32_=79.99, p=3.23e-10). There were also no differences in OmenFat between WT and KO males, but HFD males had increased OmenFat compared to LFD (F_1, 20_=46.688, p=1.22e-6) (data not shown).

**Figure 4.**
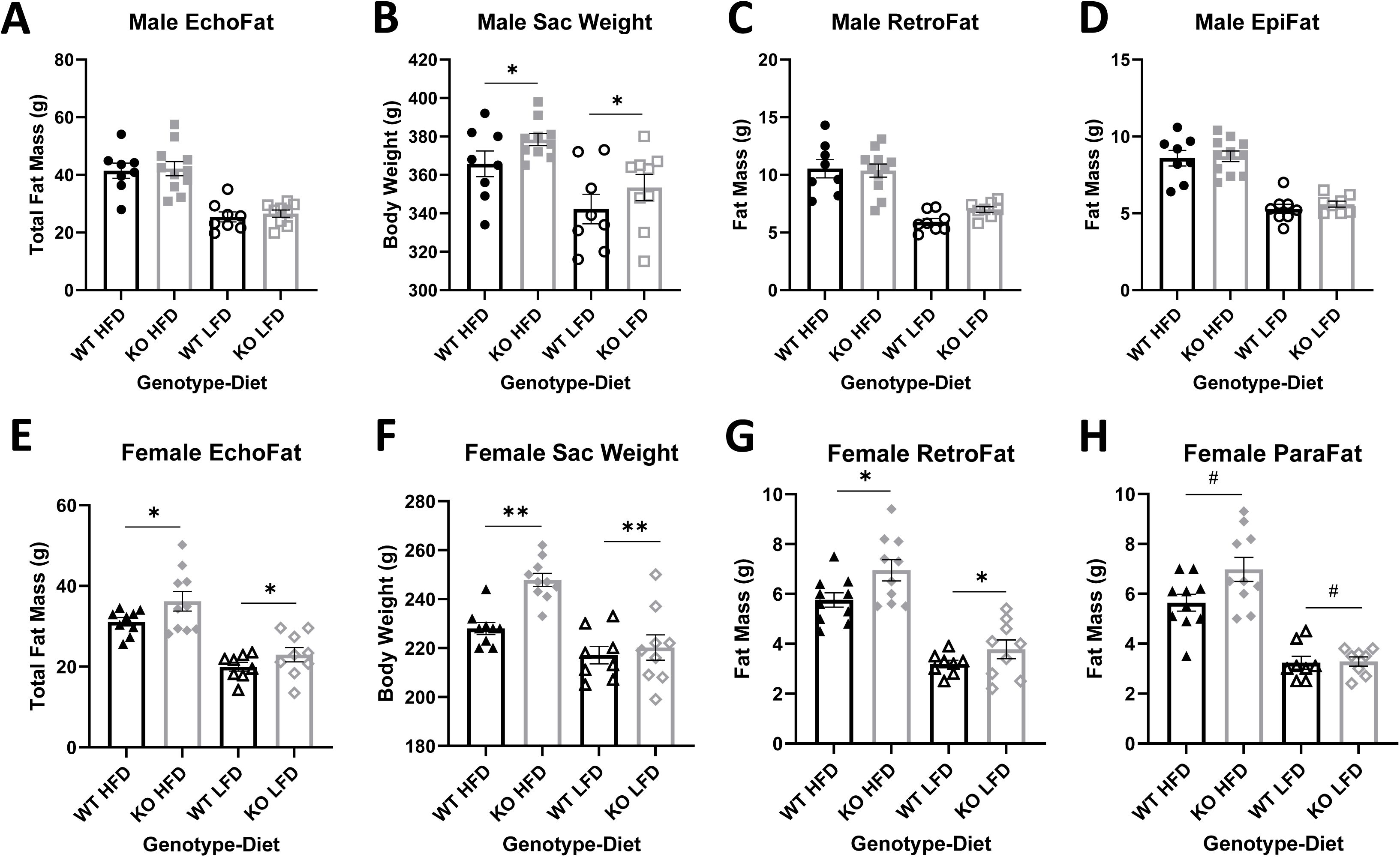
Body weight and fat mass in wild-type (WT; black) and *Krtcap3* knock-out (KO; grey) rats, for males (top) and females (bottom). After 10 weeks on diet, (A) there were no differences in total fat mass using EchoMRI analysis (EchoFat) between WT males (circle) and KO males (square), regardless of diet condition (high fat diet (HFD; filled symbols) and low fat diet (LFD; open symbols)). At euthanasia, (B) KO males were heavier than WT males, but there were no differences (C) in retroperitoneal fat (RetroFat) mass or in (D) Epididymal fat (EpiFat) mass. On the other hand, (E) KO females (diamond) on both diets had increased total fat mass using EchoMRI analysis compared to WT females (triangle), (F) increased body weight at euthanasia, (G) increased RetroFat mass, and (H) a trend toward increased parametrial fat (ParaFat) mass. Significance represents main effect of genotype at the following significance levels: ^#^p < 0.1, *p < 0.05, **p < 0.01. Diet effects were also seen as reported in the “Results” section.

### Female KO rats have increased fat mass relative to WT

After 10 weeks on diet at EchoMRI analysis, KO females had significantly greater total fat mass relative to WT females (F_1, 33_=5.32, p=0.028; **Figure 4e**). As expected, HFD females had more fat mass than LFD (F _1, 33_=48.70, p=5.59e-8). There were no differences in lean mass between WT and KO females, although HFD females did have greater lean mass than LFD females (F_1, 33_ = 4.79, p = 0.036) (**Figure S3b**). At the end of the study, KO female rats were significantly heavier than WT (F_1, 33_=9.67, p=0.0039; **Figure 4f**) and HFD females were heavier than LFD (F_1, 33_=26.58, p=1.16e-5), with no interactions between genotype and diet. In addition to increased body weight, KO females had greater RetroFat relative to WT (F_1, 34_=6.01, p=0.020; **Figure 4g**) and HFD females had greater RetroFat relative to LFD (F_1, 34_=71.90, p=6.66e-10). KO females had a trend toward increased gonadal fat relative to WT (F_1, 33_=3.38, p=0.075; **Figure 4h**) and HFD females had significantly higher gonadal fat relative to LFD (F_1, 33_=87.46, p=8.43e-11). However, there were no differences in OmenFat by genotype, although HFD females did have increased OmenFat compared to LFD (F_1, 23=_13.214, p=2e-4) (data not shown).

### Glucose tolerance is similar between WT and KO rats in both sexes

There was not a significant effect of genotype on fasting glucose in male rats (**Figure 5a**), but HFD males had increased fasting glucose relative to LFD (F_1, 22_=6.14, p=0.021). There were no differences by genotype or by diet on glucose AUC in males (**Figure 5b**).

**Figure 5.**
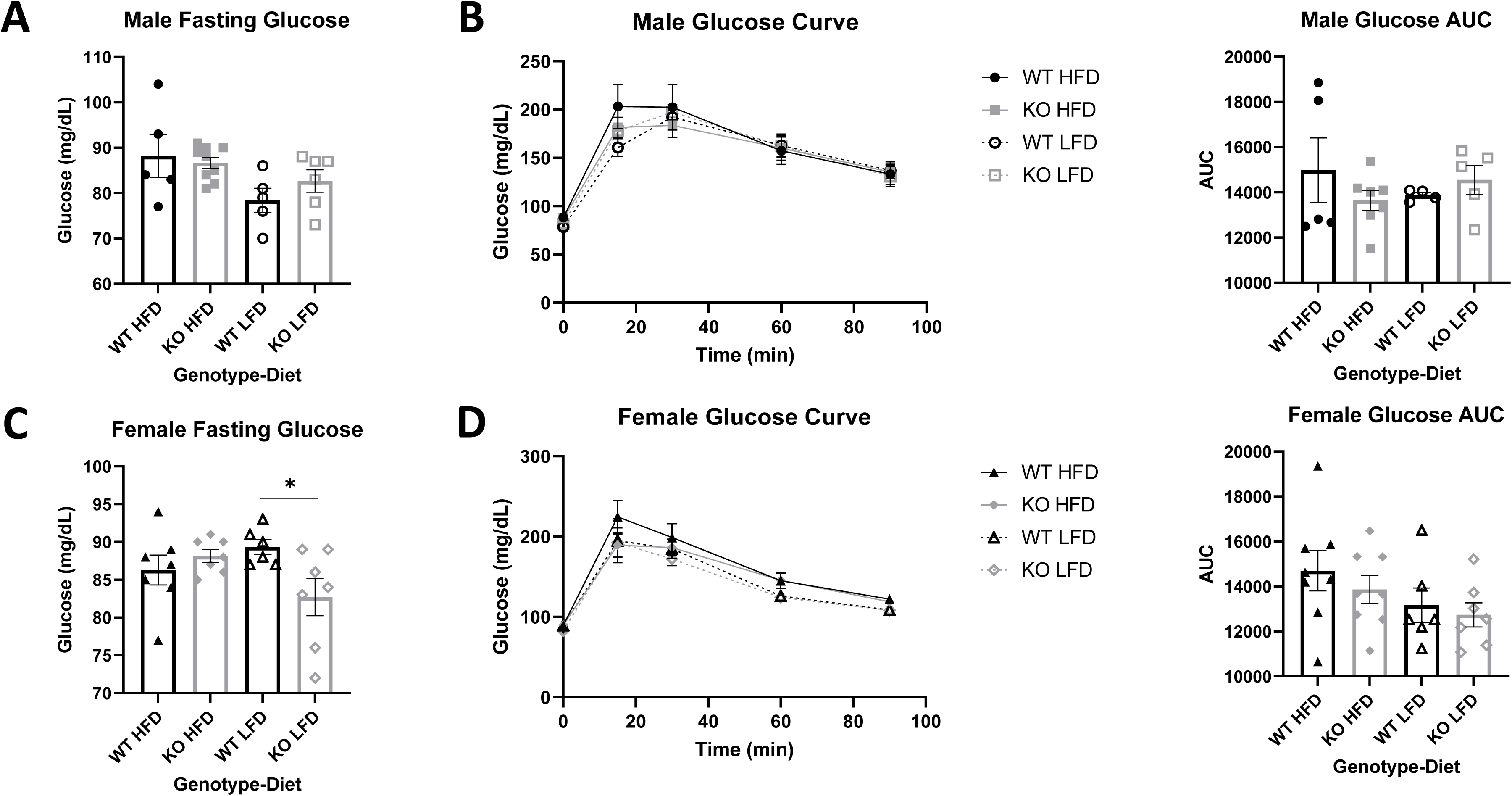
No differences in glucose response between wild-type (WT, black) and *Krtcap3* knock-out (KO, gray) rats. After 12 weeks on diet, male (top) and female (bottom) rats were fasted for 16 hours overnight in preparation for an intraperitoneal glucose tolerance test. (A) There were no differences in fasting glucose between WT males (circle) and KO males (square), neither on a high-fat diet (HFD; filled) nor a low-fat diet (LFD; empty). (B) There were also no significant differences in glucose area-under-the-curve (AUC) between WT and KO males, on either diet condition. (C) There was a significant genotype-by-diet interaction for female fasting glucose. While there was no difference in fasting glucose between WT females (triangle) and KO females (diamond) on a HFD (filled), when on a LFD (empty) KO females had a lower fasting glucose than WT females. *p < 0.05 for an effect of genotype in the LFD condition. (D) There was no significant difference in glucose AUC between WT and KO females.

There were no differences in fasting glucose by genotype or by diet in female rats, but there was a genotype by diet interaction (F_1, 24=_5.70, p=0.026) where KO LFD females had lower fasting glucose than WT LFD females (T_11=_2.51, p=0.029; **Figure 5c**), with no differences in HFD females. While there was no difference in glucose AUC by genotype (**Figure 5d**), HFD females had a slightly increased glucose AUC relative to LFD (F_1, 26_=3.38, p=0.077).

### Male KO rats are more insulin resistant than WT

There was a significant effect of genotype on fasting insulin in males (F_1, 17_=5.97, p=0.026) which was mainly driven by a genotype by diet interaction (F_1, 17_=6.82, p=0.018) where KO HFD males had increased fasting insulin relative to WT HFD males (T_11_=2.67, p=0.022) with no difference by genotype in LFD males (**Figure 6a**). As expected, HFD males had greater fasting insulin than LFD (F_1, 17_=30.43, p=3.78e-5). There was also a significant effect of genotype on HOMA-IR (F_1, 16_=12.23, p=0.003) which was also driven by a significant genotype by diet interaction (F_1, 16_=1.16, p=0.0089) where KO HFD males had higher HOMA-IR scores than WT HFD (T_10_=3.34, p=0.0074) with no difference by genotype in LFD males (**Figure 6b**). As expected, HFD males had greater HOMA-IR scores than LFD (F_1, 16_=31.59, p=3.83e-5). There was no genotype-driven effect on insulin AUC (**Figure 6c**), although HFD males had a higher insulin AUC than LFD (F_1, 20_=26.63, p=4.76e-5). There was no effect of genotype or diet on WPIC (**Figure 6d**).

**Figure 6.**
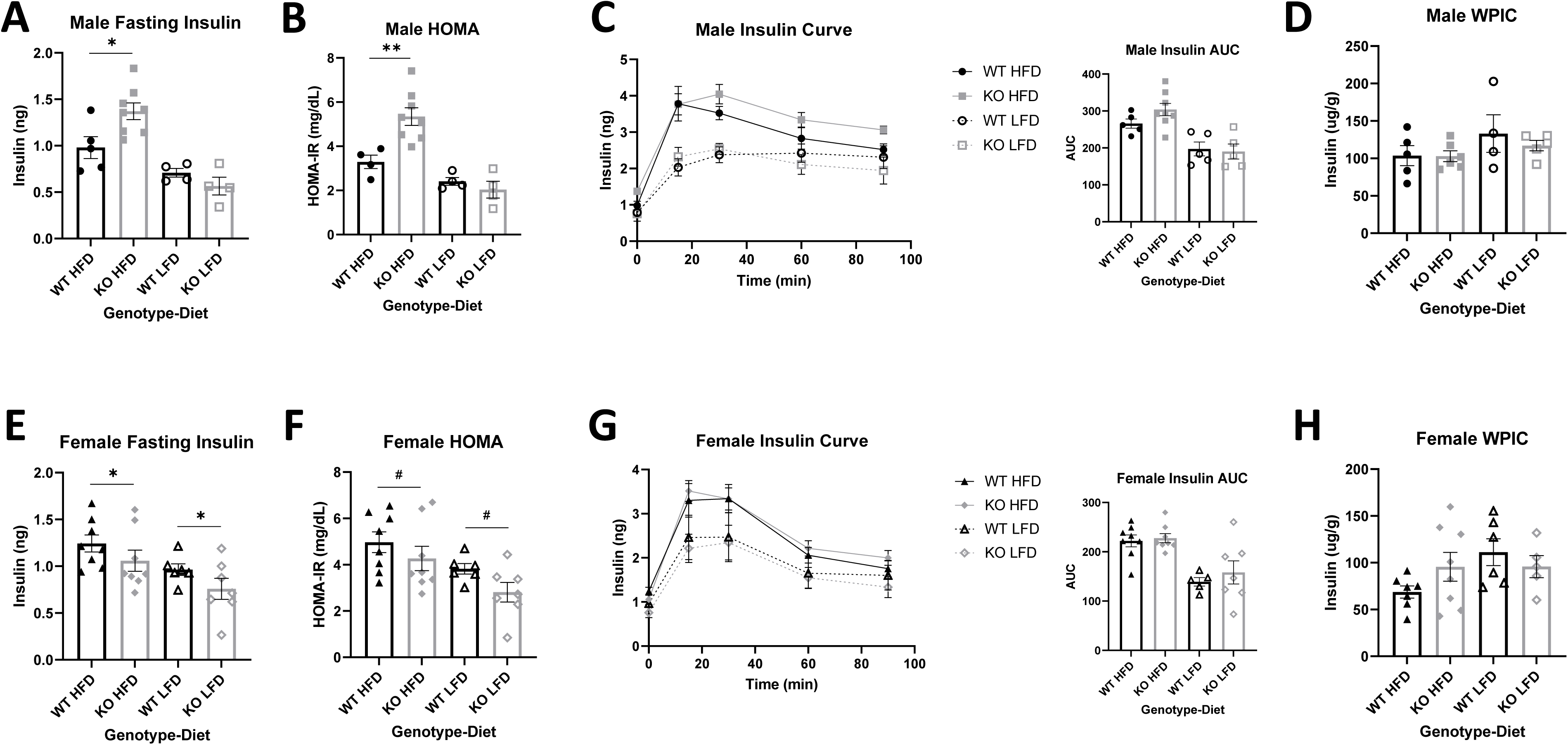
Sex differences in insulin response between wild-type (WT, black) and *Krtcap3* knock-out (KO, gray) male (top) and female (bottom) rats. Insulin was measured in serum collected from the intraperitoneal glucose tolerance test at 12 weeks on diet, and whole pancreas insulin content (WPIC) was measured per gram pancreas collected at euthanasia. (A) There was a significant genotype-by-diet interaction in male rats such that KO males (square) had a higher fasting insulin than WT males (circle) only on a high-fat diet (HFD; filled), not a low-fat diet (LFD; empty). *p < 0.05 for an effect of genotype in the HFD condition. (B) There was a genotype-by-diet interaction for homeostatic model assessment of insulin resistance (HOMA-IR) in male rats such that KO males had a higher HOMA-IR score than WT males only when on a HFD. **p < 0.01 for an effect of genotype in the HFD condition. (C) There were no significant differences in insulin area-under-the-curve (AUC) between WT and KO males. (D) There were no significant differences in WPIC between WT and KO males. (E) KO females (diamond) on both diets had a lower fasting insulin than WT females (triangle). *p < 0.05 for a main effect of genotype. (F) KO females had a trend toward a lower HOMA-IR score compared to WT females. ^#^p < 0.1 for a main effect of genotype. (G) There were no differences, however, in insulin AUC between WT and KO females and (H) there were no significant differences in WPIC between WT and KO females of either diet. Diet effects are reported in the “Results” section.

### Female KO rats are more insulin sensitive than WT

There was a significant effect of genotype on fasting insulin in female rats, but in the opposite direction of the males, with KO females having decreased fasting insulin relative to WT (F_1, 26_=4.31, p=0.048; **Figure 6e**). Similar to males, HFD females had increased fasting insulin relative to LFD (F_1, 26_=8.56, p=0.0071). There was a nearly significant main effect of genotype on HOMA-IR (F_1, 26_=4.12, p=0.053; **Figure 6f**) with increased HOMA-IR in WT females relative to KO, and as expected increased HOMA-IR in HFD females relative to LFD (F_1, 26_=8.83, p=0.0063). As with males, while there was no difference in insulin AUC by genotype (**Figure 6g**), HFD females did have a greater insulin AUC than LFD (F_1, 26_=9.04, p=0.0058). There was also no effect of genotype or of diet for WPIC (**Figure 6h**).

### No differences in serum or liver lipids between WT and KO males

There were no significant differences by genotype or by diet in serum triglycerides in male rats (**Figure 7a**). While there was no effect of genotype on serum total cholesterol (**Figure 7b**), HFD males had lower cholesterol than LFD (F_1, 33_=4.46, p=0.042). Similarly, there were no differences in liver triglycerides between WT and KO males (**Figure 7c**), but HFD males had lower liver triglycerides than LFD (F_1, 33_=12.46, p=0.0013). There were no differences by genotype or by diet in liver cholesterol (**Figure 7d**).

**Figure 7.**
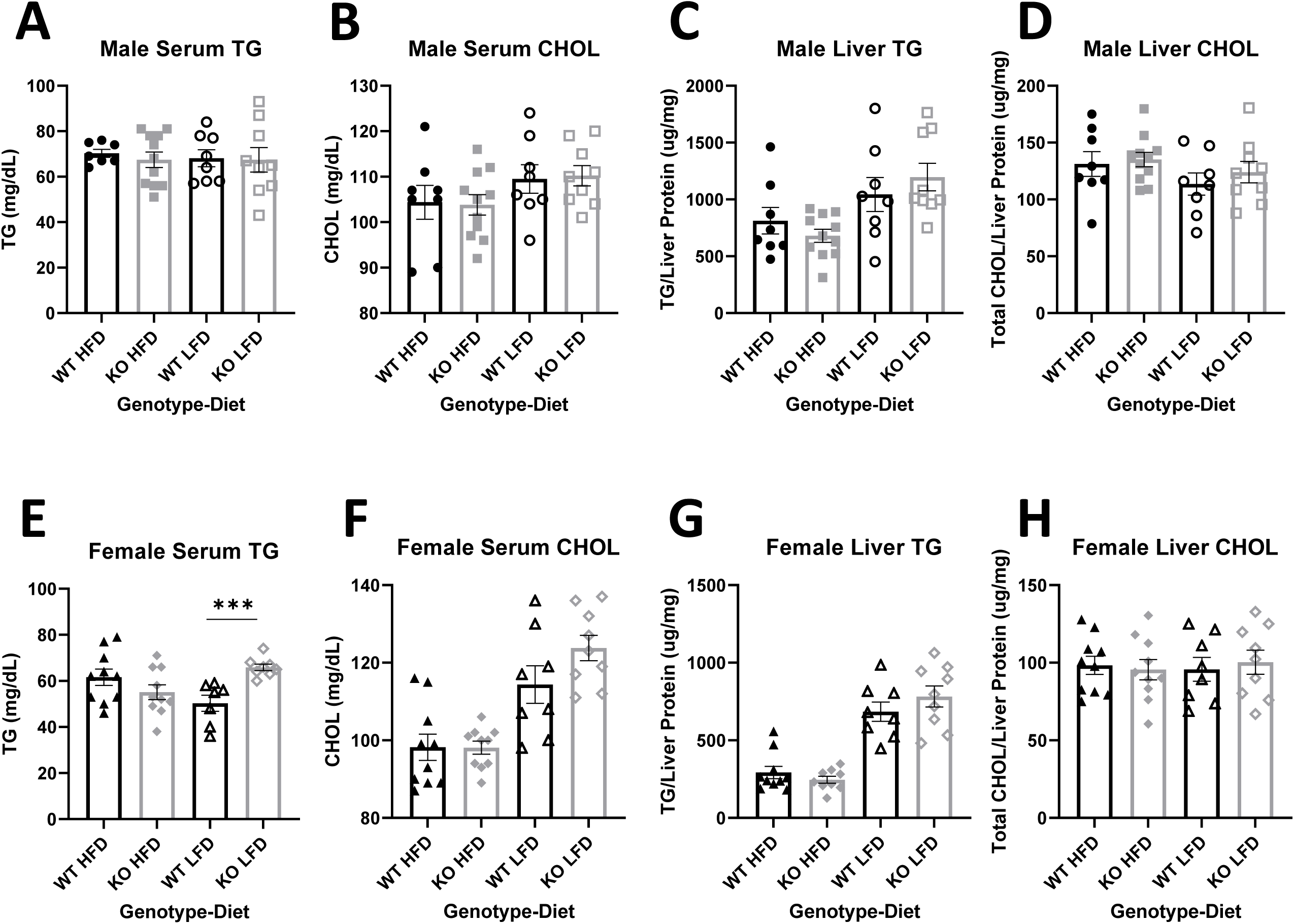
No differences in liver metabolic health between wild-type (WT; black) and *Krtcap3* knock-out (KO; gray). Triglycerides (TG) and cholesterol (CHOL) were measured in serum collected from male (top) and female (bottom) rats. (A) There was no difference between WT males (circle) and KO males (square) on a high-fat diet (HFD; filled) or a low-fat diet (LFD; empty) for (A) serum TG or (B) serum CHOL. TG and CHOL were also measured in liver tissue of male rats, with no differences between WT and KO males for (C) liver TG or (D) liver CHOL. (E) There was a genotype-by-diet interaction in serum TG in female rats, such that while there was no difference between WT (triangle) and KO females (diamond) on a HFD, KO females (diamond) had increased serum TG compared to WT (triangle) on a LFD. ***p < 0.001 for an effect of genotype in the LFD condition. (F) There were no differences in serum CHOL between WT and KO females. There were no differences in (G) liver TG nor (H) liver CHOL between WT and KO females. Diet effects are reported in the “Results” section.

### No pathological differences in serum or liver lipids between WT and KO females

There were no differences by genotype or by diet in serum triglycerides for female rats but there was a genotype by diet interaction (F_1, 31=_12.01, p=0.0016) where only KO LFD females had higher triglycerides relative to WT LFD females (T_13=_14.35, p=0.00079; **Figure 7e**), with no difference in HFD females. Genotype did not affect serum total cholesterol (**Figure 7f**), but HFD females had lower cholesterol than LFD (F_1, 34_=39.75, p=3.47e-7). Similarly, there were no differences in liver triglycerides by genotype (**Figure 7g**), but HFD females had lower liver triglycerides relative to LFD (F_1, 33_=96.77, p=2.44e-11). As with the males, there were no differences by genotype or diet in liver cholesterol (**Figure 7h**).

### No differences in adipocyte size or number between WT and KO HFD females

As expected, there were few very large adipocytes in both WT and KO rats, with a significant effect of adipocyte bin size (F_29, 269_=321, p=2e-16), but surprisingly there were no differences by genotype in the adipocyte area frequency distributions (**Figure 8a**). Representative images demonstrate this lack of difference (**Figure 8b**).

**Figure 8.**
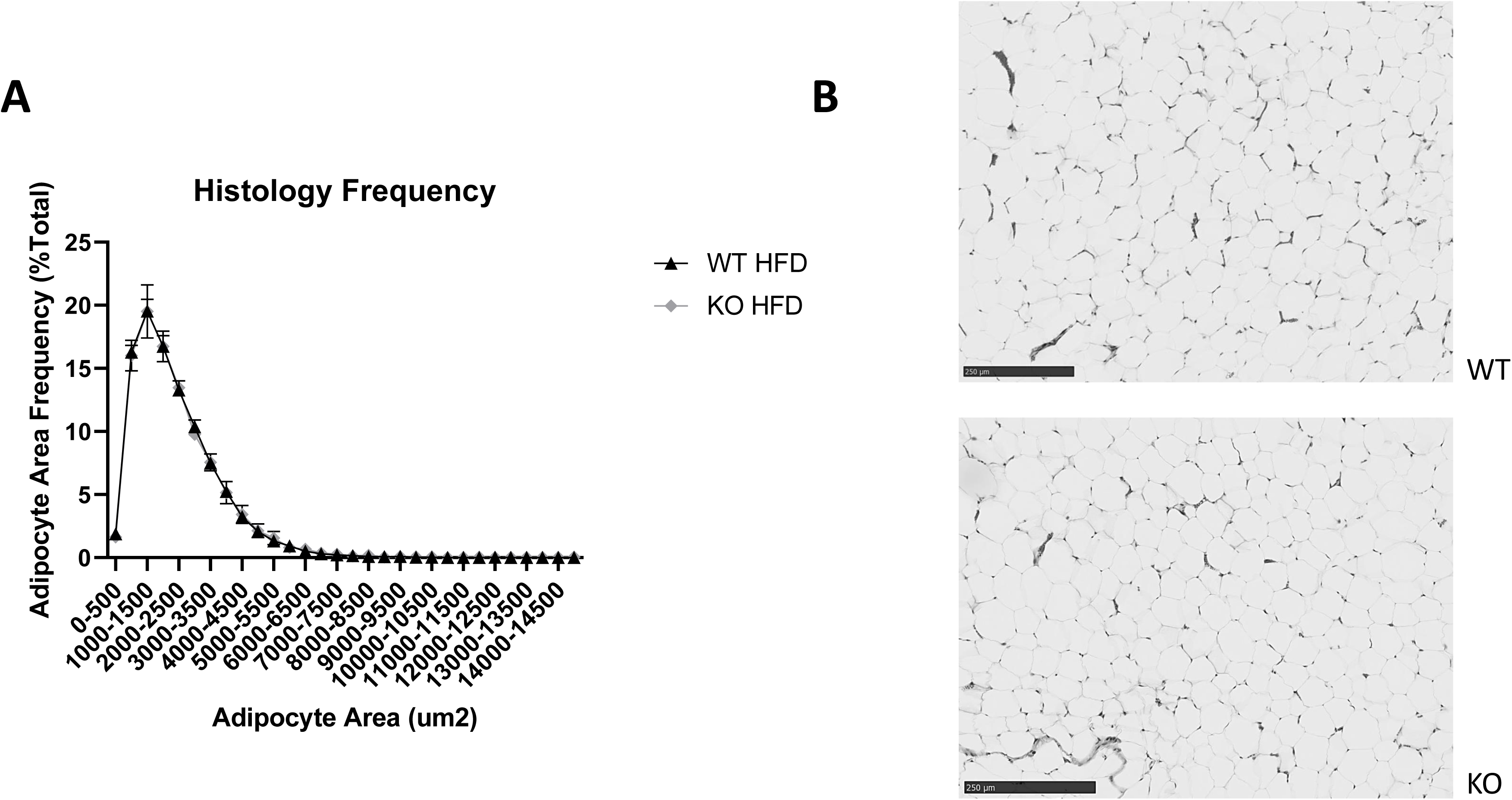
No difference in adipocyte size or number between wild-type (WT; black filled triangle) and *Krtcap3* knock-out (KO; gray filled diamond) female rats. Sections of retroperitoneal fat (RetroFat) were taken from representative WT and KO females on a high-fat diet (HFD) and stained for hematoxylin and eosin. (A) There were no differences in the frequency distributions of adipocyte area between WT and KO females, as shown by (B) representative adipocytes between WT (top) and KO (bottom) rats. Images were re-colored to grayscale and sharpened by 25%.

## Discussion

We demonstrate that *Krtcap3* plays a role in body weight in both sexes, but that only females exhibit increased eating and adiposity. Despite increased food intake, body weight, and fat mass, KO females had increased insulin sensitivity and no differences in adipocyte size or frequency, suggesting a metabolically healthy phenotype. In contrast, while we saw increased body weight in KO male rats, we were unable to detect significant differences in eating behavior or fat mass compared to WT males. These data validate *Krtcap3* as a novel gene for adiposity, implicate its role in food intake and insulin sensitivity, and demonstrate important sex differences.

*Krtcap3* has previously been identified as a candidate adiposity gene in both humans and rats (7, 11). Despite these findings, little is known about the underlying function of *Krtcap3*, with previous work showing a role in sheep wool production and some human cancers (14, 15, 16, 17), and essentially no validation or functional studies for its role in adiposity. Expression patterns of *Krtcap3* show moderately high levels in male and female sex organs and the gastrointestinal (GI) tract in humans and/or rats (https://gtexportal.org; https://ncbi.nlm.nih.gov), which may help inform the current findings, where we saw important sex and food intake differences.

At six weeks of age, KO females were heavier than WT controls, and these differences became more pronounced when the rats aged, particularly when on a HFD. That said, KO females on both diets had increased fat mass relative to WT. We propose that increased food intake drives the adiposity phenotype in the KO females. This is not surprising, as findings from human GWAS for BMI are enriched in the central nervous system and may play a role in satiety signaling (4, 18, 19, 20). Although the *Krtcap3*-KO exhibited increased adiposity, these rats did not show worsened metabolic health. Specifically, there were not any pathological genotype-driven differences in glucose or lipid metabolism, but KO female rats were more insulin sensitive compared to WT counterparts. Furthermore, while adipocyte hypertrophy is associated with human obesity (21), there were no differences in adipocyte area between WT and KO females. These data point to *Krtcap3* being a gene that increases body weight and fat mass in female rats without the associated co-morbidities. Interestingly, recent human GWAS have identified genetic variants that increase body fat percentage but are protective against metabolic complications (22). This supports the idea that *Krtcap3* may promote weight gain without diminishing metabolic health, at least in females.

In male rats, we found that KO rats were heavier than WT controls at six weeks of age, and that this body weight difference was maintained throughout the study. Unexpectedly, we did not see increased adiposity in the male KO compared to WT rats, nor were there significant eating differences. The body weight difference is most likely explained by the slight increase in lean mass in KO males compared to WT males. The lack of an adiposity difference was surprising, as *Krtcap3* was originally identified as a candidate gene in male rats (7). Despite not seeing an adiposity phenotype, we found that KO HFD males had hyperinsulinemia and insulin resistance compared to WT controls, with no differences in LFD males. Similar to females, there were no genotype-driven differences in lipid metabolism.

Sex differences in obesity have been well-established in rodents and humans (23, 24, 25, 26) and may lend support to the striking sex differences seen here. The differences we see may be explained by *Krtcap3* having a larger effect size in females relative to males, which is supported by literature on sex-specific obesity loci, where loci explain more variance in women than men (27, 28). Future replication studies to increase the *n* may allow us to see an effect in males. The second explanation is that *Krtcap3* may interact with gonadal hormones to cause sex differences (24). Sex steroid hormones are known to play a role in food intake, fat deposition, and even diabetes susceptibility (24, 26, 29). Estrogens in particular can suppress appetite, alter fat deposition, and protect against insulin resistance (26, 29). An interaction between *Krtcap3* and sex hormones may explain the different adiposity and insulin phenotypes between male and female rats in the present study. Future studies will explore these hypotheses.

As the original study identified *Krtcap3* expression in liver (7), we anticipated that KO rats would have increased serum and liver lipids. Surprisingly, there were no pathological genotype-driven differences in these measures in either male or female rats. These data suggest that while liver *Krtcap3* expression is associated with increased fat mass, its role in the liver is likely not driving the increased adiposity. The food intake phenotype points toward a potential role for *Krtcap3* in satiety signaling, but *Krtcap3* has low expression in the brain regions associated with food intake (30). Because *Krtcap3* is highly expressed in the gut, another possibility is that *Krtcap3* may play a role in the gut-brain axis to affect food intake (31). Future work is necessary to investigate these hypotheses.

This work validates *Krtcap3* as a novel gene for adiposity. We show that female rats exhibit increased body weight, food intake, and fat mass with improved metabolic health. In contrast, male rats exhibit increased body weight with worsened metabolic health and no change in fat mass or food intake. Further work will explore the sex differences and investigate both where and how *Krtcap3* is acting to impact adiposity.

## Acknowledgments

The authors thank Dr. Kylie Kavanagh and lab for allowing use of her Visiopharm software, Matt Davis for his help with liver lipid profiling, and Rebecca Schilling and Allison Zappa at the MCW Genotyping Core for their assistance in genotyping the rats.

## Supplementary Methods

### Animals

To develop the *Krtcap3*-KO (WKY-Krtcap3^em3Mcwi^) we injected a CRISPR targeting the sequence GGGACTTGCGCTGATCCTGG into Wistar-Kyoto (WKY/NCrl; RGD_1358112) rat embryos, and the resulting mutation was a net 2 base pair deletion in exon 2 (**Figure 1a**). This led to a frameshift mutation and early termination of the Krtcap3 protein. Founder animals were genotyped by the Cel-1 assay [32] and confirmed by Sanger sequencing. Founders were then backcrossed to the parental WKY strain and subsequent litters at the Medical College of Wisconsin (MCW) were genotyped by fluorescent genotyping.

We compared *Krtcap3* expression in liver and adipose tissue between available inbred founder strains (ACI, BN, BUF, F344, M520, and WKY) (**Figure S1b**). RNA was extracted by Trizol and expression was measured using real-time quantitative PCR (rt-qPCR). *GAPDH* (**Table S1**) was used as the housekeeping gene and fold change of *Krtcap3* (**Table S1**) transcript was calculated by the following equation:

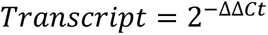

Where ΔCt was the difference between the crossing threshold (Ct) of *Krtcap3* and *GAPDH*, and ΔΔCt the difference between each sample ΔCt and the average ACI ΔCt.

Experimental wild-type (WT) and knock-out (KO) rats were genotyped one of two ways. The first was fluorescent based fragment analysis at MCW using the ABI 3730 capillary sequencer, followed by analysis in Genemapper software, as mentioned above. The second method, at Wake Forest School of Medicine (WFSoM), was to amplify *Krtcap3* over the mutation site, and then digest the samples with the restriction enzyme MboI. WT samples would be digested into a smaller strand, while KO samples, which lacked the MboI binding site due to the mutation, would remain the same size as the original PCR product. When run out on a 1.5% DNA agarose gel, WT and KO samples were distinguished by their banding pattern. The same primer for *Krtcap3* amplification was used for genotyping at MCW and at WFSoM (**Table S1**).

To verify that the knock-out reduced *Krtcap3* expression, we collected liver tissue from adolescent WT and KO rats (n = 3 per genotype). RNA was extracted by Trizol and expression was measured using rt-qPCR. *β-actin* (**Table S1**) was used as the housekeeping gene, and fold change of *Krtcap3* transcript was calculated by the equation given above, but where ΔCt was the difference between the crossing threshold (Ct) of *Krtcap3* and *β-actin*, and ΔΔCt the difference between each sample ΔCt and the average WT ΔCt.

### Statistical Analysis

All data were assessed visually by a boxplot for potential outliers. When n > 6, Grubbs’ test for outliers was performed, and if p < 0.05, the suspected outlier was removed. In cases with a lower n the interquartile range method was used to detect outliers. Values falling outside the upper and lower bounds were removed. Normality was assessed by a Shapiro-Wilk test and by visually inspecting a Q-Q plot. Homogeneity of variance was also assessed by the Bartlett test.

## Supplementary Tables

**Table S1.**
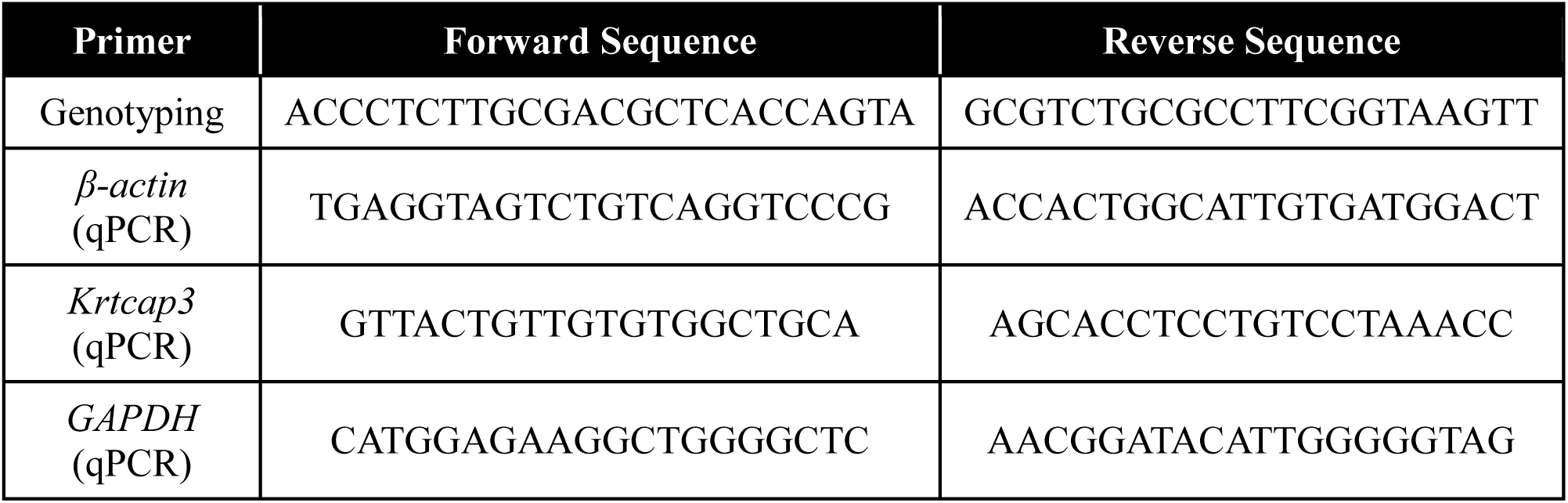
Primer sequences. 5’ → 3’

**Table S2.** Dietary composition in %kcal. LFD, low-fat diet; HFD, high-fat diet.

|  | **LFD (D12450J)** | **HFD (D12492)** |
| --- | --- | --- |
| Protein | 20 | 20 |
| Carbohydrate | 70 | 20 |
| Fat (Lard + Soybean Oil) | 10 | 60 |
| Total kcal/g | 3.82 | 5.21 |

**Table S3.** Number of rats per experimental procedure. Male (M), Female (F), wild-type (WT), knock-out (KO), high-fat diet (HFD), low-fat diet (LFD). Experimental procedures include wean weight (Wean), six-week body weight (6-week), EchoMRI analysis (EchoMRI), intraperitoneal glucose tolerance test (IPGTT), euthanasia (Sac), and whole pancreas insulin content (WPIC).

| **Sex-Genotype-Diet** | **Wean** | **6-week** | **EchoMRI** | **IPGTT** | **Sac** | **WPIC** |
| --- | --- | --- | --- | --- | --- | --- |
| **M WT HFD** | 14 | 16 | 8 | 5 | 8 | 4 |
| **M WT LFD** |  |  | 8 | 5 | 8 | 4 |
| **M KO HFD** | 18 | 22 | 11 | 9 | 11 | 6 |
| **M KO LFD** |  |  | 9 | 6 | 9 | 5 |
| **F WT HFD** | 16 | 18 | 10 | 8 | 10 | 8 |
| **F WT LFD** |  |  | 8 | 6 | 8 | 6 |
| **F KO HFD** | 17 | 19 | 10 | 8 | 10 | 8 |
| **F KO LFD** |  |  | 9 | 7 | 9 | 5 |

**Table S4.** Data transformations. TG, triglycerides; ParaFat, parametrial fat; CHOL, cholesterol. Phenotypes not listed had a normal distribution and variance and did not need to be transformed.

| Sex | Phenotype | Type of Transformation | Normality/Variance |
| --- | --- | --- | --- |
| Male | Fasting Glucose | Reciprocal root | Normality & Variance |
|  | Glucose AUC | Natural log | Normality |
|  | Liver TG | Natural log | Normality |
| Female | ParaFat | Natural log | Normality |
| | Fasting Glucose | $\wedge 4$ | Normality |
|  | Serum CHOL | Natural log | Normality |
|  | Liver TG | Natural log | Normality |

## Supplementary Figure Legends

**Figure S1.**
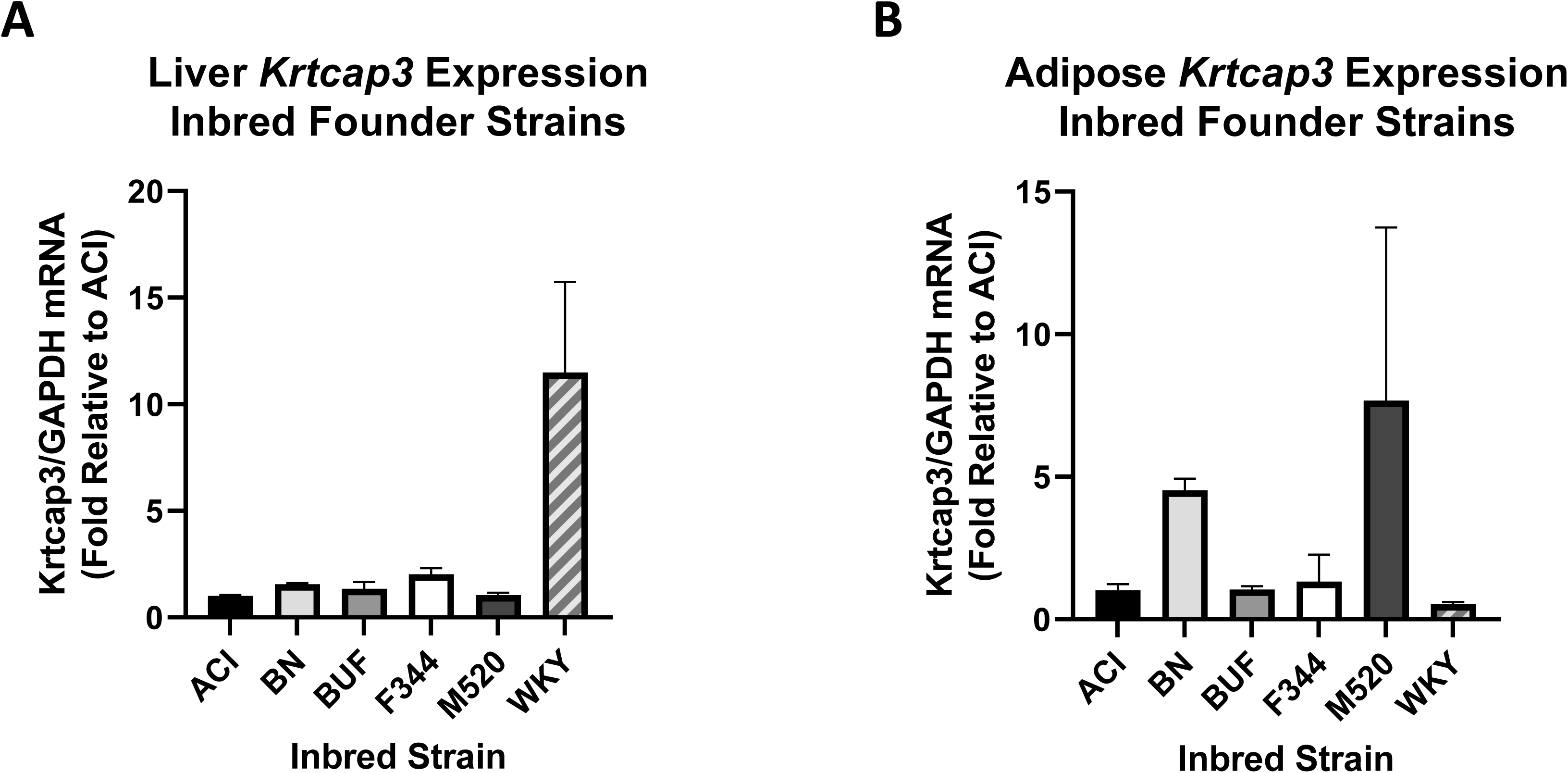
Comparison of *Krtcap3* expression between six of the heterogeneous stock rat inbred founder strains. (A) RNA was extracted from liver tissue of available inbred founder strains to compare *Krtcap3* expression. Available strains are ACI (black), BN (light gray), BUF (medium gray), F344 (white), M520 (dark gray), and WKY (striped gray). Fold change was calculated relative to ACI *Krtcap3* expression. (B) RNA was extracted from retroperitoneal adipose tissue of the same inbred founder strains to compare *Krtcap3* expression. Fold change was calculated relative to ACI *Krtcpa3* expression.

**Figure S2.**
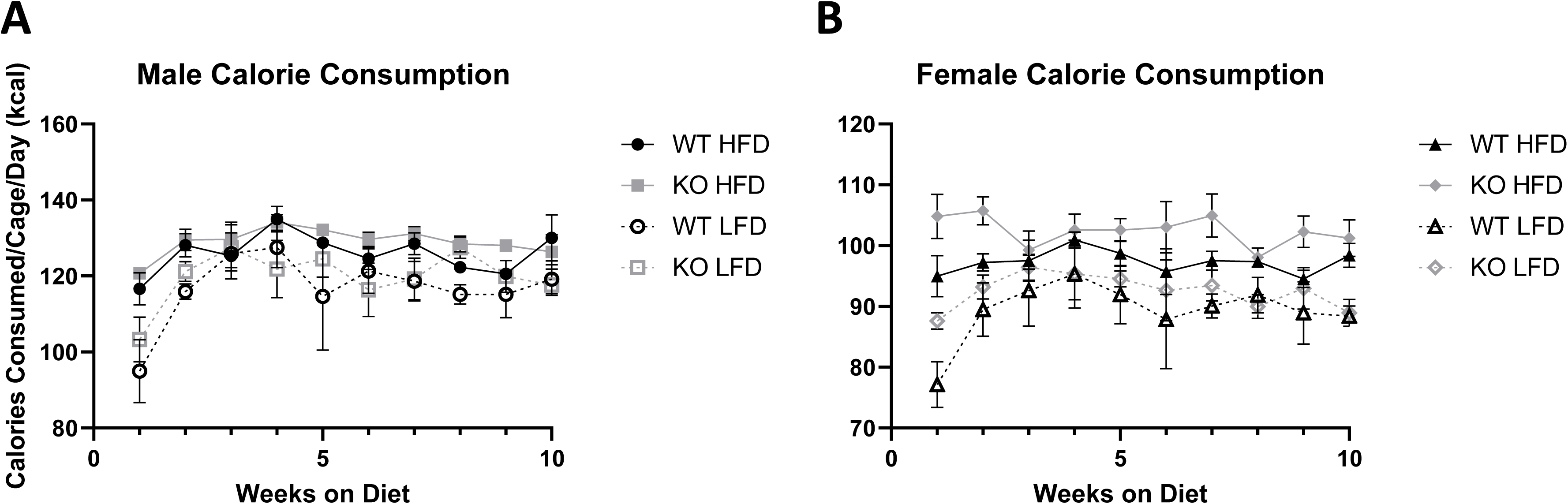
High-fat diet (HFD; filled) rats consumed more calories than low-fat diet (LFD; empty) rats. (A) While there were no differences in calories consumed between wild-type (WT; black circle) and *Krtcap3* knock-out males (KO; gray square), in both genotypes HFD males consumed more calories than LFD males. (B) There were differences in eating behavior by genotype in females, resulting in KO females (gray diamond) consuming more calories than WT females (black triangle); HFD females also consumed more calories than LFD females.

**Figure S3.**
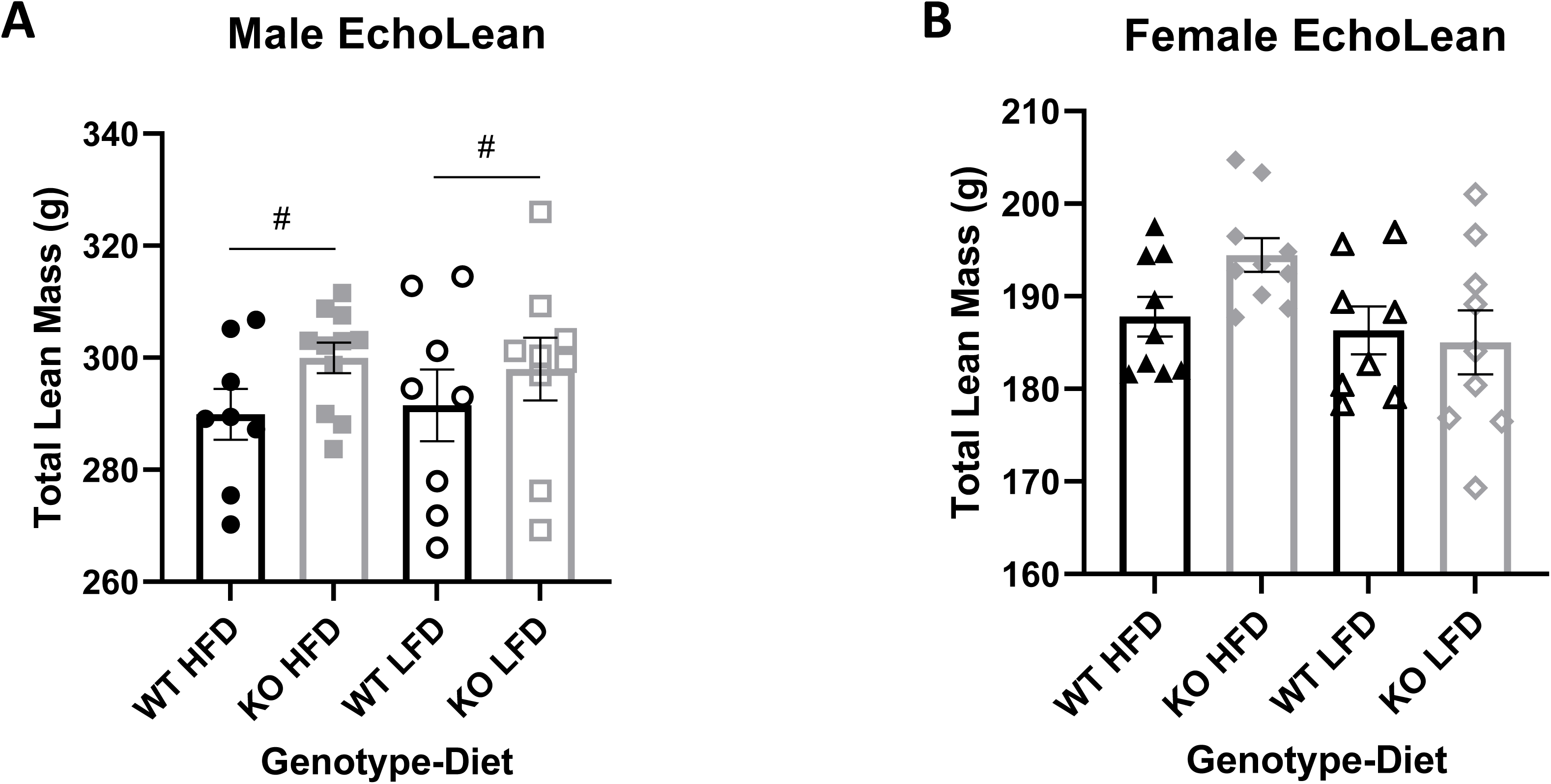
Lean mass in wild-type (WT; black) and *Krtcap3* knock-out (KO; gray) rats. After 10 weeks on diet, (A) there were slight differences in lean mass between WT males (circle) and KO males (square), but no differences between males on a high-fat diet (HFD; filled) or a low-fat diet (LFD; empty). (B) There were no differences in lean mass between WT females (triangle) and KO females (diamond) in either diet condition. Significance represents main effect of genotype at the following significance level: ^#^p < 0.1. Diet effects in females are reported in the “Results” section.

